# Spatiotemporally controlled Myosin relocalization and internal pressure cause biased cortical extension to generate sibling cell size asymmetry

**DOI:** 10.1101/311852

**Authors:** Tri Thanh Pham, Arnaud Monnard, Jonne Helenius, Erik Lund, Nicole Lee, Daniel J. Müller, Clemens Cabernard

## Abstract

Metazoan cells can generate unequal sized sibling cells during cell division. This form of asymmetric cell division depends on spindle geometry and Myosin distribution but the underlying mechanics are unclear. Here, we use atomic force microscopy and live cell imaging to elucidate the biophysical forces involved in the establishment of physical asymmetry in Drosophila neural stem cells. We show that the force driving initial apical membrane expansion is provided by hydrostatic pressure, peaking shortly after anaphase onset, and enabled by a relieve of actomyosin contractile tension on the apical cell cortex. The subsequent increase in contractile forces at the cleavage furrow, combined with the relocalization of basally located Myosin results in basal membrane extension and sustained apical expansion. We propose that spatiotemporally controlled actomyosin contractile tension and hydrostatic pressure enables stereotypic biased membrane expansion to generate sibling cell size asymmetry.

## Introduction

Sibling cell size asymmetry - here also called physical asymmetry - refers to the formation of unequal sized cells during cell division. Metazoan cells tightly regulate the mechanisms controlling symmetric or asymmetric physical cell divisions but the underlying mechanisms and physiological roles are still unclear (Cabernard, 2017; Roubinet and Cabernard, 2014). The generation of equal sized sibling cells can be regulated through spindle geometry. For instance, changing spindle position or the location of the metaphase plate of symmetrically dividing cultured human cells can induce physical asymmetric cell divisions (Kiyomitsu and Cheeseman, 2013; Tan et al., 2015). *Drosophila* male germline stem cells, generating an equal sized gonialblast and a self-renewed germline stem cell (Spradling et al., 2011), control spindle symmetry with the centrosomal-associated microtubule depolymerizing kinesin Klp10A (Kinesin 13 family); *klp10A* mutants contain unequal sized centrosomes, resulting in the formation of asymmetric spindles and unequal sized cell divisions (Chen et al., 2016). Similarly, ER localized Kif2A, a microtubule depolymerase, induces spindle asymmetry and positioning in ascidians (Costache et al., 2017). Since the anaphase spindle is the primary determinant for the positioning of the cleavage furrow (D’Avino et al., 2015; Glotzer, 2017; Green et al., 2012; Rappaport, 1986), spindle geometry can at least partially explain physical asymmetry.

Whereas these cell types regulate the formation of equal sized siblings, neuroblasts in *C. elegans* and *Drosophila melanogaster* divide asymmetrically by size. *C. elegans* Q neuroblasts give rise to a large neuron and a small apoptotic cell and changing the intrinsically controlled sibling cell size ratio affects subsequent cell fate and behavior (Ou et al., 2010). *Drosophila* neuroblasts (NBs; neural stem cells) form a large self-renewed neuroblast and a small differentiating ganglion mother cell (GMC) (Gallaud et al., 2017; Homem and Knoblich, 2012). Although changes in cell polarity affect spindle geometry and sibling cell size in fly neuroblasts (Albertson and Doe, 2003; Cabernard et al., 2009; Cai et al., 2003), recent findings suggest that neuroblasts in *C. elegans* and *Drosophila melanogaster* regulate sibling cell size asymmetry through asymmetric localization of Non-muscle Myosin II (Myosin hereafter) (Cabernard et al., 2010; Connell et al., 2011; Ou et al., 2010). Drosophila neuroblasts are intrinsically polarized (Gallaud et al., 2017; Homem and Knoblich, 2012) and utilize the apically localized polarity protein Partner of Inscuteable (Pins; LGN/AGS3 in vertebrates) to induce asymmetric Myosin localization (Cabernard et al., 2010; Connell et al., 2011). Pins induces an apical enrichment of activated Myosin through Drosophila Rho Kinase (Rok) and Protein Kinase N (Pkn) (Tsankova et al., 2017). At anaphase onset, Myosin relocalizes to the cleavage furrow through (1) a basally-directed and – with ~ 1 minute delay – (2) an apically-directed cortical Myosin flow. The molecular mechanisms triggering apical – basal cortical Myosin flow onset are not entirely clear but involve apical Pins and potentially other neuroblast intrinsic polarity cues. On the basal neuroblast cortex, spindle-dependent cues induce an apically directed cortical Myosin flow to flow to the cleavage furrow. The correct timing of these Myosin flows – regulated by polarity and spindle cues – is instrumental in establishing biased Myosin localization and sibling cell size asymmetry in fly neuroblasts (Roth et al., 2015; Roubinet et al., 2017; Tsankova et al., 2017).

Although biased Myosin localization provides a framework for the generation of unequal sized sibling cells (Connell et al., 2011) it is currently unknown what ultimately drives biased membrane expansion. Myosin localization dynamics, controlling actomyosin contractile tension, could spatiotemporally affect cortical stiffness but other biophysical forces could contribute to the establishment of physical asymmetry. Here, we use atomic force microscopy to measure dynamic changes in cell stiffness and cell pressure, combined with live cell imaging and genetic manipulations in asymmetrically dividing neuroblasts. We found that physical asymmetry is formed by two sequential events: (1) internal pressure initiates apical expansion due to a Myosin-dependent softening of the apical neuroblast cortex. (2) Actomyosin constriction at the basally-shifted cleavage furrow subsequently initiates basal expansion and maintains apical membrane expansion. Thus, spatiotemporally coordinated Myosin relocalization combined with hydrostatic pressure and cleavage furrow constriction enables biased membrane extension and the establishment of stereotypic sibling cell size asymmetry.

## Results

### A cell intrinsic stiffness asymmetry precedes the formation of the cleavage furrow

Cell shape changes are largely controlled by changes in mechanical stress and tension at the cell surface (Clark et al., 2015). During physical asymmetric cell division, cortical proteins are subject to precise spatiotemporal control, but how this impacts cell surface tension to allow for complex cell shape changes is incompletely understood (Figure 1A). To this end, we set out to measure the dynamic changes in cell stiffness of asymmetrically dividing neuroblasts with Atomic Force Microscopy (AFM). Larval brain neuroblasts *in vivo* are surrounded by cortex glia on the apical, and ganglion mother cells (GMCs) and differentiating neurons on the basal side, preventing us from directly measuring changes in stiffness on the neuroblast surface. We thus established primary neuroblast cultures from third instar larval brains so that the AFM tip can directly probe the neuroblast surface. Cultured larval brain neuroblasts show normal polarization and divide with comparable cell cycle timing to neuroblasts *in vivo* (Supplemental Figure 1A-C and (Berger et al., 2012; Cabernard et al., 2009)). We used a rounded AFM tip with a 300 nm diameter to measure neuroblast stiffness - a measure for the resistance of the cell surface to an applied external force – on ~ 20 positions along the apical – basal division axis. These measurements, performed every 30s on neuroblasts expressing Sqh::GFP (Royou et al., 2002) (labelling Myosin’s regulatory light chain) and GFP-tagged centrosomin (cnn::GFP; labelling centrosomes (Zhang and Megraw, 2007)) were averaged over multiple cells (n = 25) and binned in 5 regions (apical, subapical, middle, sub-basal and basal) (Figure 1B). Sqh::GFP and Cnn::GFP were used to (1) determine the cell cycle stage and (2) map the position of where the AFM tip touched the neuroblast surface. Since we found that cell elongation rates usually peaked at around 90s after anaphase onset, we used cnn::GFP to detect the cell elongation onset and thereby to determine the mitotic stage; these cell elongation rate measurements allowed us to determine anaphase onset with ~ 30s accuracy (Supplemental Figure 1D,E). Unreliable AFM measurements, visible in the force curves, were excluded from the analysis (Supplemental Figure 1F-H and methods).

**Figure 1:**
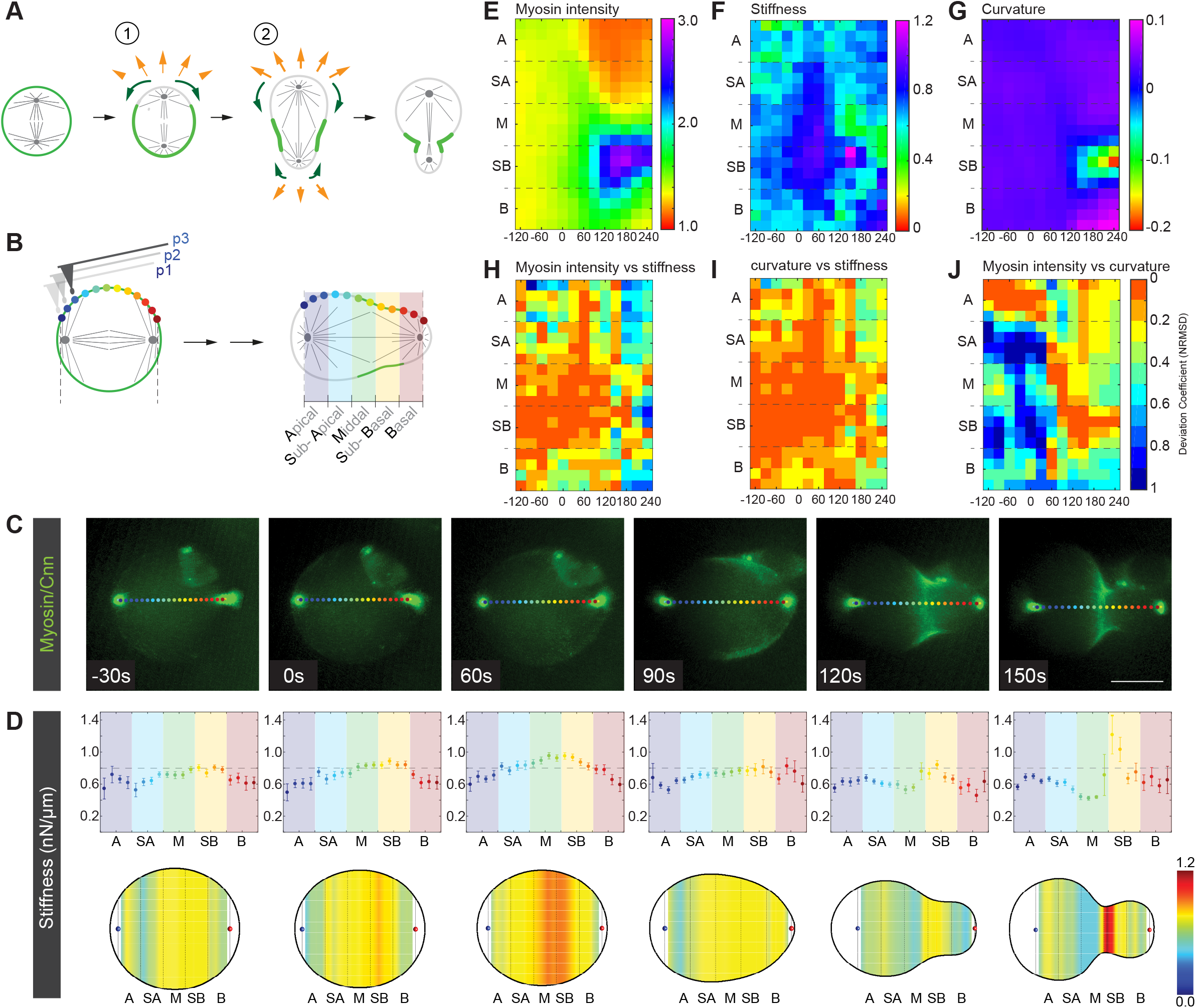
Cortical stiffness only partially correlates with Myosin localization and curvature. (**A**) Wild type neuroblasts undergo biased membrane expansion (orange arrows) concomitant to spatiotemporally controlled Myosin relocalization (green arrows). Apical Myosin flows (green arrows) towards the cleavage furrow prior to the onset of an apically directed Myosin flow (green arrows). The underlying force for biased membrane expansion is incompletely understood. (**B**) Schematic representation showing cortical stiffness measurement points along the cell cortex (colored circles) throughout mitosis. Measurements were binned into five cortical regions along the apical-basal neuroblast axis. (**C**) Representative image sequence showing a wild type neuroblast expressing Sqh::GFP (Myosin; green) and the centrosome marker Cnn::GFP (bright green dots) throughout mitosis. Positions where AFM measurements were performed are labelled with colored circles. (**D**) Distribution of mean cortical stiffness and standard error of the mean (top row, n = 25) in all regions along the division axis throughout mitosis in reference to anaphase onset (0 s). The bottom row shows the corresponding heat map for cortical stiffness. (**E**) Mean Myosin intensity (n = 19) at all sub-binned regions for wild type neuroblasts; time axis is relative to anaphase onset. Mean stiffness (n = 25) and mean curvature (dimensional unit is *μ*m^−1^, n = 19) are shown in (**F**) and (**G**), respectively. Deviation coefficients (see methods) were plotted to correlate Myosin intensity with cortical stiffness (**H**), curvature with stiffness (**I**) and Myosin intensity with curvature (**J**). Scale bar: 5 *μ*m.

Our AFM measurements revealed that cell stiffness is mostly uniform before anaphase onset ranging from 0.4-0.8 nN/*μ*m. At anaphase onset and 60s thereafter (“0s” marks anaphase onset in Figure 1C), we observed a slight decrease in stiffness at both cell poles and a noticeable increase in stiffness in the Mid and Sub-basal region, reaching almost 1.0 nN/*μ*m. Interestingly, this increase in stiffness appeared off cell center and closer to the basal cell cortex in a region of the prospective cleavage furrow. Subsequently, this asymmetric stiffness increase disappeared and stiffness dropped over most of the cell cortex despite an apparent lateral increase in Myosin intensity ~90s after anaphase onset. Also, stiffness predominantly dropped on the apical cell pole. Stiffness increased again in late anaphase, predominantly in the cleavage furrow region. These measurements revealed a surprisingly dynamic pattern of stiffness changes during asymmetric cell division.

### Neuroblast stiffness is a combination of actomyosin contractile tension and other biophysical parameters

Previously, it was suggested that cortical relaxation at the poles was responsible for biased membrane expansion during physical asymmetric cell division (Connell et al., 2011). Cortical relaxation could be induced through Myosin relocalization, prompting us to correlate stiffness changes with Myosin relocalization dynamics. Since the wide field data were not sufficiently reliable to extract Myosin intensity, we imaged third instar neuroblasts expressing Sqh::GFP with spinning disc microscopy and correlated the resulting intensity and curvature profiles with AFM stiffness data by calculating the relative change between Myosin and curvature, and Myosin and stiffness. As reported previously (Roubinet et al., 2017), apical Myosin intensity started to decrease at anaphase onset, albeit stiffness increased again apically. In the mid and sub-basal region, both Myosin intensity and stiffness increased. Similarly, high Myosin intensity was visible at the forming cleavage furrow later in anaphase, concomitant with detectable changes in cell surface curvature (Figure 1E, F). The most noticeable curvature changes became apparent in the furrow region from 120s after anaphase onset onwards (Figure 1E,F,G). The resulting deviation coefficients revealed that Myosin intensity and curvature strongly correlate on the early (−120 – 30s) apical neuroblast cortex and later in the cleavage furrow region (120s - 240s); the shift in Myosin intensity from the apical cell cortex towards the furrow region – previously described as a cortical flow (Roubinet et al., 2017) – was accompanied with a shift in curvature changes. Until 120s after anaphase onset, stiffness correlated best with Myosin intensity in the cleavage furrow region. However, it is noticeable that in many cortical regions, Myosin intensity showed a poor correlation with either curvature and/or stiffness (Figure 1H-J).

These measurements revealed a surprisingly dynamic pattern of local stiffness changes which only partially correlate with Myosin localization and cell shape changes. From these data we conclude the following: (1) stiffness asymmetries exist along the apical-basal neuroblasts axis prior to furrow ingression. (2) The accumulation in Myosin at the cleavage furrow correlates well with changes in curvature. (3) Local changes in stiffness only partially correlate with local Myosin localization. We hypothesize that the local accumulation of Myosin filaments directly or indirectly affects cell surface properties in cortical regions with low Myosin filament concentration. Furthermore, the data suggests that the registered neuroblast stiffness is a combination of Myosin activity as well as other biophysical parameters.

### Hydrostatic neuroblast pressure drops in early anaphase

Since cells increase their hydrostatic pressure during mitosis (Stewart et al., 2011), we wondered whether changes in neuroblast stiffness can be attributed to changes in hydrostatic pressure. We used a parallel plate assay to measure rounding force by pressing a wedge onto cultured neuroblasts expressing the membrane marker PH::GFP (Mavrakis et al., 2009; Ramanathan et al., 2015; Stewart et al., 2013) (see also methods and Figure 2A). Rounding force gradually increased during mitosis before dropping sharply shortly after anaphase onset (Figure 2B). From these measurements, we used two methods to calculate the corresponding hydrostatic pressure: (1) we measured the surface area in contact with the AFM wedge for each time point and divided the registered force by this value. (2) We used the Young-Laplace formula (YONEDA, 1980; 1964)(see also methods) to obtain the contact area before anaphase; at these stages, Young-Laplace calculations are very precise since cells are predominantly spherical prior to elongation in anaphase. However, our contact area measurements were much higher than the calculated Young-Laplace surface area, probably due to optical aberrations. We calculated a correction factor and applied it to the detected contact area to extract hydrostatic pressure at all time points (Figure 2C, D). These measurements show that cultured neuroblasts increased their intracellular hydrostatic pressure up to 105 Pa (mean P = 103; SD = +/- 6.08; n = 13) 1 minute after anaphase onset. However, 2 minutes after anaphase onset, hydrostatic pressure has already dropped by 20% compared to its peak value. Once division is completed, neuroblasts contain a significantly lower hydrostatic pressure (mean P = 10.92; SD = +/-3.72; n =13) compared to the onset of mitosis (Figure 2E, F). These data show that the changes in hydrostatic pressure closely matches neuroblast stiffness dynamics, suggesting that hydrostatic pressure is a major contributor to neuroblast stiffness.

**Figure 2:**
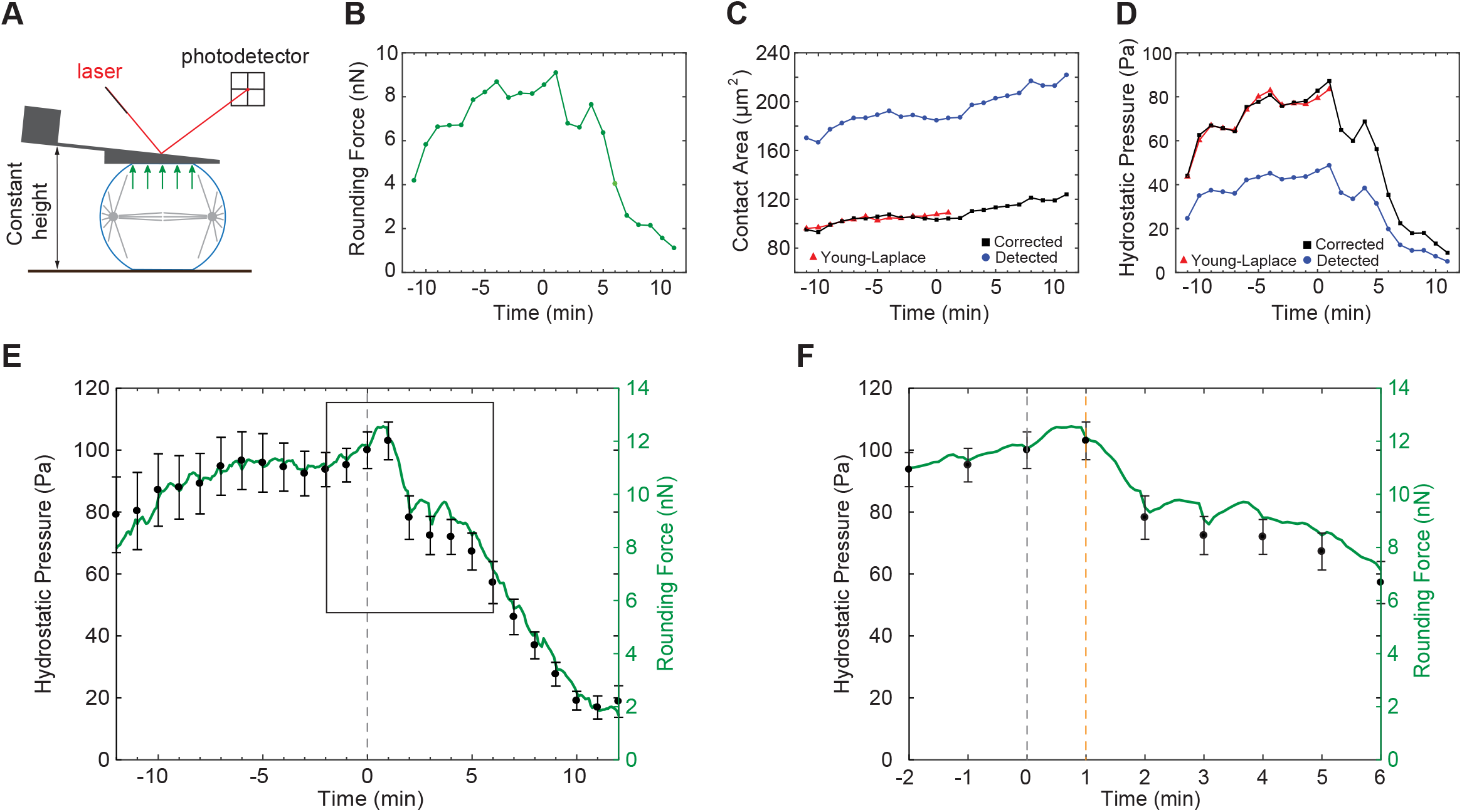
Hydrostatic pressure increases during mitosis, peaking right after anaphase onset. (**A**) Schematic representation of a parallel plate assay (constant height assay) used to measure hydrostatic pressure throughout mitosis. (**B**) Rounding force detected by the AFM throughout mitosis for a representative cell. The time axis is relative to anaphase onset (0 s). (**C**) Contact area was determined by two independent methods. (1) Theoretical contact area (red) calculated using the uniform tension model is very accurate, however it is only valid as long as cells are spherical. (2) Contact area detected by the total cell area that are in contact with the AFM wedge using fluorescent signal (blue). This method can be used at all cell cycle stages, but it tends to overestimate the contact area due to the point spread function of the imaging system. A correction factor was applied to contact area measurements based on Young-Laplace calculations for spherical cells (**D**). Hydrostatic pressure for the chosen representative cell was plotted per Young-Laplace (red), the detected contact area (blue) and the corrected contact area (black). (**E**) Resulting mean rounding force and mean hydrostatic pressure (error bars represent standard error of the mean, n=13) for the corrected contact area throughout mitosis for wild type neuroblasts. The boxed measurements are shown enlarged in (**F**). The grey vertical line refers to anaphase onset; the orange vertical line highlights the drop in hydrostatic pressure.

### Hydrostatic pressure is coordinated with Myosin relocalization to drive biased membrane expansion

Previously, we showed that Myosin relocalization dynamics strongly correlate with physical asymmetry (Connell et al., 2011; Roubinet et al., 2017; Tsankova et al., 2017) but the force underlying biased membrane expansion remained unexplained. Neuroblast membrane extension could be driven by (1) furrow constriction, displacing fluid and cytoplasmic material, (2) internal pressure or (3) a combination of both (Figure 3A). We thus analyzed how dynamic changes in neuroblast pressure during mitosis correlate with Myosin relocalization, constriction and biased membrane expansion. To this end, we imaged neuroblasts expressing Sqh::GFP and the spindle marker cherry::Jupiter and generated kymographs by drawing a line along the apical-basal axis and the cleavage furrow (Figure 3B-D & Supplemental Figure 2A, B). We used these kymographs to quantify (1) the extend of apical and basal cortical expansion in relation to anaphase onset and cleavage furrow ingression, (2) constriction and expansion rates and (3) Myosin intensity at the apical, basal and furrow cortex (Figure 3E-G). These measurements showed that wild type neuroblasts always started to expand shortly after anaphase onset on the apical neuroblast cortex first, followed by furrow diameter reduction. The expansion of the basal cell cortex occurred almost at the same time as furrow constriction was detectable (Figure 3H). Prior to constriction, neuroblasts expanded by more than 1 *μ*m on the apical cortex but no expansion was detected basally. Once constriction started, expansion was measureable on both the apical and basal neuroblast cortex (Figure 3I, J). These data suggest that (1) apical expansion is primarily driven by hydrostatic pressure and (2) sustained apical and all basal expansion is driven by furrow constriction. Furthermore, biased cortex expansion could correlate with Myosin relocalization dynamics. Indeed, apical expansion occurred shortly after anaphase onset, coinciding with a drop in apical Myosin intensity and high internal pressure. The onset of basal cortex expansion - ~ 90s after anaphase onset – coincided with a drop in basal Myosin, low hydrostatic pressure but an increase in Myosin intensity at the cleavage furrow (Figure 3K).

**Figure 3:**
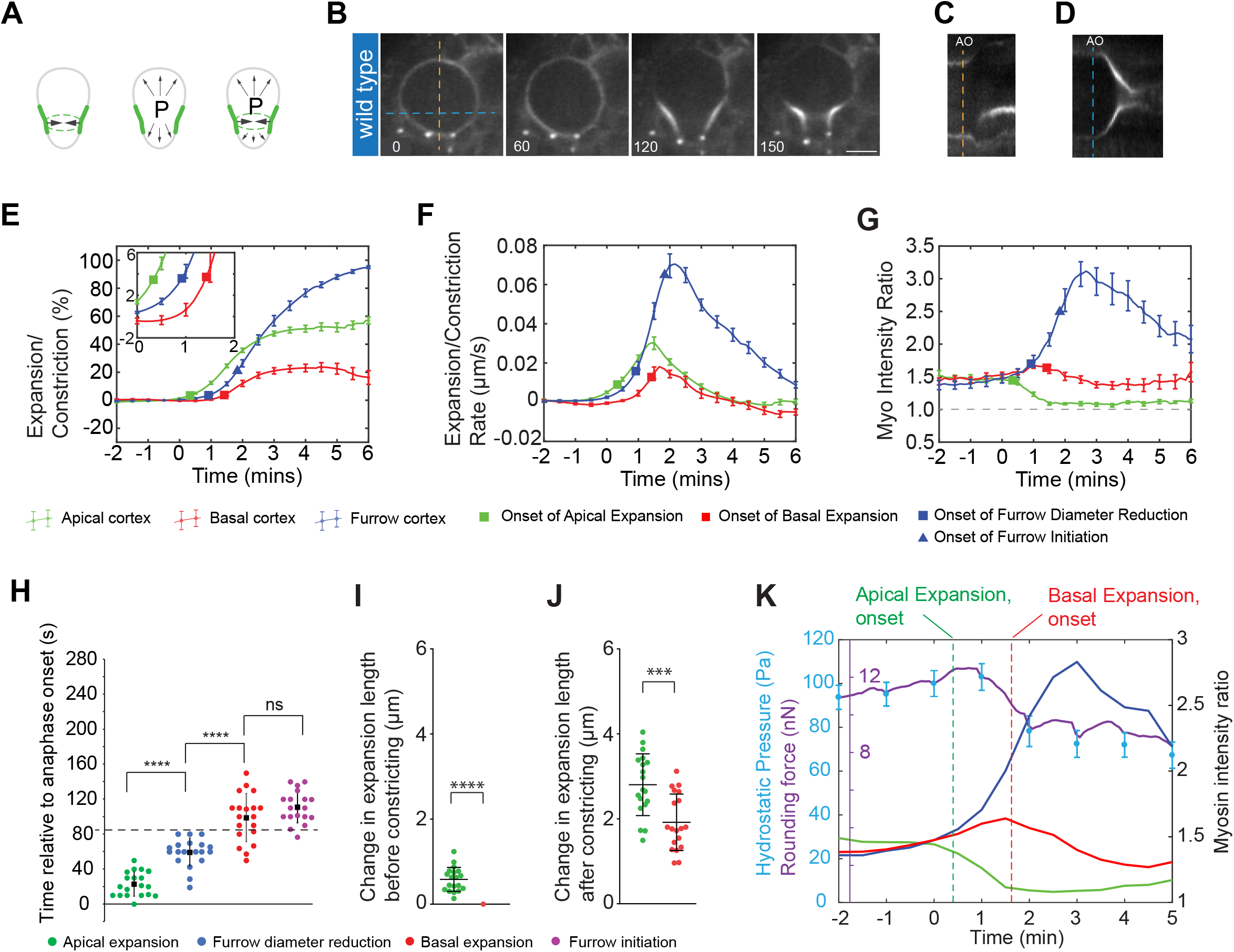
Internal hydrostatic pressure and Myosin’s relocalization dynamics drive asymmetric cortical expansion in fly neuroblasts. (**A**) Biased membrane expansion could be driven by furrow constriction, intracellular hydrostatic pressure or a combination of both. (**B**) Representative image sequence showing a wild type neuroblast expressing Sqh::GFP (white). Kymographs obtained along the apical-basal axis (orange dotted line) are shown in (**C**) or the furrow region (blue dotted line) in (**D**). Mean change in the expansion/constriction length (**E**), rate of change (**F**) and Myosin intensity (**G**) are plotted for the apical cortex (green), the basal cortex (red) and the furrow site (blue). Vertical bars refer to standard error of the mean (n=19). The time axis is relative to anaphase onset (0 s). (**H**) Scatter plot showing the onset of apical expansion (green), onset of furrow diameter reduction (blue), onset of basal expansion (red) and onset of furrow initiation (when the curvature first changes from a straight line to an inward bending curve; purple). Changes in expansion length for both apical (green) and basal (red) cortex before and after furrow constriction are shown in (**I**) and (**J**), respectively. (**K**) Graph showing the mean rounding force (purple), mean hydrostatic pressure (cyan circles with error bars), mean Myosin intensity at the apical cortex (green), mean Myosin intensity at the basal cortex (red) and mean Myosin intensity at the furrow site cortex (blue). The green and red dashed lines represent the mean onset of apical and basal expansion, respectively. Scale bar: 5 *μ*m. Asterisk denote statistical significance, derived from unpaired t-tests: *; p ≤ 0.05, **; p ≤ 0.01, ***; p ≤ 0.001, ****; P ≤ 0.0001, n.s.; not significant.

Taken together, these data suggest that apical membrane expansion is driven by high internal hydrostatic pressure and permitted through decreasing actomyosin contractile tension on the apical cortex. Basal membrane expansion however is primarily driven by an increase in cleavage furrow constriction and enabled by a lowering of actomyosin contractile tension on the basal cell cortex.

### Spatiotemporal control of actomyosin contractile tension affects expansion dynamics and sibling cell size asymmetry

Next, we tested how global or local modulations in actomyosin contractile tension affected biased membrane extension dynamics and sibling cell size asymmetry. *pins* mutant neuroblasts have been shown to alter Myosin relocalization dynamics, clearing Myosin from both the apical and basal cell cortex simultaneously (Cabernard et al., 2010; Roubinet et al., 2017; Tsankova et al., 2017). In contrast to wild type, apical and basal extension occurred to the same extent already prior to furrow constriction (Figure 4A, H, F, G & Supplemental Figure 2C, D). Since it was shown that cell rounding underlies a balance between actomyosin contractile tension and hydrostatic pressure (Stewart et al., 2011), we hypothesized that equal cortical expansion should be primarily driven by intracellular pressure, consistent with the expansion dynamics of *pins* mutant neurolasts. Thus, lowering actomyosin contractile tension by reducing the amount of activated cortical Myosin should also lower intracellular hydrostatic pressure and thereby altering expansion dynamics. Indeed, adding the Rho kinase inhibitor Y-27632 to wild type neuroblasts resulted in significantly lower cortical Myosin intensity; complete Rok inhibition showed no difference between cortical and cytoplasmic signal, whereas partial inhibition still contained lowered cortical Myosin levels (Figure 4B, C & Supplemental Figure 2E-H). Strong Rok inhibition prevented all apical and basal membrane extension and constriction (Figure 4B and Supplemental Figure 2E) but partial inhibition of Rok allowed apical and basal membranes to expand at the onset of furrow diameter reduction. However, most expansion occurred predominantly after furrow constriction set in (Figure 4B, C, F, G, I). Thus, lowering actomyosin contractile tension shifted the initial pressure driven expansion towards constriction-driven expansion. To test how an increase in actomyosin contractile tension affects cortical expansion dynamics, we trapped Myosin at the cell cortex by coexpressing Sqh::GFP together with the membrane tethered nanobody (pUAST-CAAX::vhhGFP4); vhhGFP4 has a high affinity for GFP (Saerens et al., 2005)), thereby retaining activated Myosin uniformly at the cell cortex. We found that similar to Rok inhibitor treated neuroblasts, expansion predominantly occurred during furrow ingression (Figure 4D, F, G, J). Finally, we attempted to bias Myosin relocalization dynamics by removing the mitotic spindle using colcemid, a condition delaying basal Myosin relocalization while still permitting normal apical clearing (Roth et al., 2015; Roubinet et al., 2017). The removal of the mitotic spindle also increased hydrostatic pressure in human cells (Stewart et al., 2011). Although the lack of a mitotic spindle prevented us from determining anaphase onset, we found that expansion only occurred apically prior to constriction, followed by a late constriction driven basal expansion event. Since Myosin was retained on the basal cortex, the basal cortex initially retracted and we measured the extend of expansion following this initial retraction only (Figure 4E, F, G, K).

**Figure 4:**
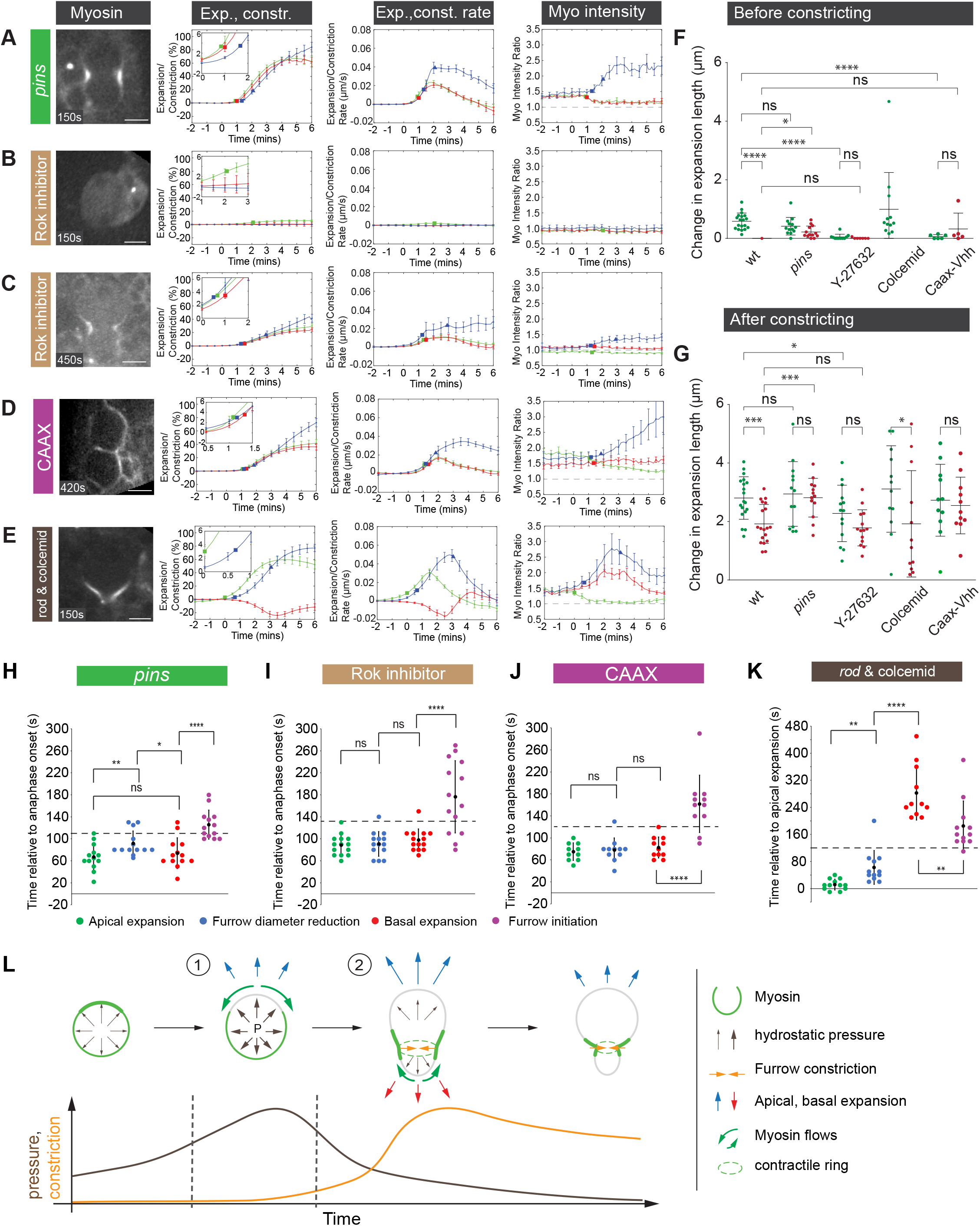
Physical asymmetric cell division is a two-step process driven by internal pressure and spatiotemporally controlled Myosin relocalization. Representative anaphase images, change in expansion/constriction length, expansion/constriction rate and Myosin intensity shown for (**A**) *pins* (n=13), (**B**) complete Rok inactivation (n=7), (**C**) partial Rok inactivation (n=14), (**D**) CAAX::vhhGFP4 expressing (n=11) and (**E**) colcemid treated, *rod* mutant (n=12) neuroblasts expressing Sqh::GFP (white). Measurements are shown for the apical cortex (green), the basal cortex (red) and the furrow site (blue). Time axis is relative to anaphase onset (0 s). Scatter plots showing the change in expansion length for the apical (green) and basal (red) cortex before (**F**) and after (**G**) constriction. Scatter plots showing the onset of apical expansion (green), furrow diameter reduction (blue), basal expansion (red) and furrow initiation (purple) for (**H**) *pins,* (**I**) partial Rok inactivation, (**J**) CAAX::vhhGFP4 expressing and (**K**) *rod* mutant neuroblasts exposed to colcemid. (**L**) Model. Spatiotemporally regulated Myosin relocalization permits initial internal pressure-driven apical expansion. Subsequently, due to dissipation of internal pressure, sustained apical and subsequent basal expansion is driven by actomyosin-dependent furrow constriction. See text for details. Scale bars: 5 *μ*m. Asterisk denote statistical significance, derived from unpaired t-tests: *; p ≤ 0.05, **; p ≤ 0.01, ***; p ≤ 0.001, ****; p ≤ 0.0001, n.s.; not significant.

Taken together, these data suggest that manipulating Myosin’s activity and localization dynamics not only affects biased cortical expansion but also determines to what extent it is driven by intracellular pressure and constriction. We conclude that intracellular pressure can only drive membrane expansion if actomyosin contractile tension is permissive. In the absence of hydrostatic pressure, or if actomyosin contractile tension is maintained, membrane expansion is primarily driven by cortical constriction.

## Discussion

Sibling cell size asymmetry occurs in multiple cell types and organisms but the underlying mechanisms are diverse and incompletely understood (Cabernard, 2017; Roubinet and Cabernard, 2014). Previously, we showed that biased cortical expansion correlates with Myosin’s distribution (Connell et al., 2011). Whereas apical Myosin relocalization dynamics are controlled by the apical polarity protein Pins, through biased localization of Rok and Pkn (Cabernard et al., 2010; Tsankova et al., 2017) basal Myosin relocalization is controlled by the spindle-dependent pathway (Roth et al., 2015; Roubinet et al., 2017). Both apical and basal Myosin relocalization occurs through cortical Myosin flow (Roubinet et al., 2017). However, the forces underlying biased membrane extension remained unclear. Here, we characterized changes in cell stiffness in correlation with Myosin localization. These measurements revealed that neuroblasts increase their surface stiffness during mitosis, peaking shortly after anaphase onset. The increase in cell stiffness is predominantly attributed to an increase in intracellular pressure and only partially to local accumulation of Myosin. Our data further revealed that apical neuroblast expansion occurs at a time point when intracellular pressure is high, enabling the relaxed apical cortex to expand. Since basal Myosin relocalization occurs with ~ 1 minute delay (Roubinet et al., 2017), pressure is insufficient to overcome basal actomyosin contractile tension until relocalization starts. Basal membrane expansion is thus driven by the constricting cleavage furrow (Figure 4L). Manipulating the spatiotemporal contractile Myosin tension further changed the relative contribution of pressure and constriction to membrane expansion.

Our stiffness measurements further revealed a characteristic tension asymmetry, marking the position of the prospective cleavage furrow. This finding is consistent with our previous measurements, showing an increase in actomyosin accumulation prior to measureable furrowing (Roubinet et al., 2017). We hypothesize that this local increase in activated actomyosin could induce furrow constriction and thus the displacement of cytoplasmic material, contributing to biased membrane expansion, in particular in the absence of high intracellular pressure.

Taken together, our data provides an intuitive model (Figure 4L), explaining the establishment of physical asymmetry. Neuroblasts utilize a combination of cell cycle and polarity regulated Myosin relocalization dynamics while concomitantly increasing intracellular pressure for biased membrane extension. Previously it was proposed that local modulations in cortical tension could allow cells to alter their shape (Stewart et al., 2011). Our study provides experimental evidence for this model under physiological conditions and could be potentially relevant to other invertebrate and vertebrate cells alike (Shin et al., 2014). In the future, it will be interesting to learn whether biased membrane extension is accompanied by asymmetric membrane addition or an unfolding of membrane stores. Similarly, the mechanisms regulating hydrostatic pressure increase during the cell cycle remain to be defined.

## Acknowledgements

We thank members of the Cabernard lab for helpful discussions. This work was supported by the Swiss National Science Foundation (SNSF; PP00P3_159318 to CC and 310030B 160255 to D.J.M), the National Institute of Health (NIH; 1R01GM126029-01, CC) and start-up funds from the University of Washington. T.T.P was supported with a Systems X Transition Postdoc Fellowship (TPdF: SXFSIO_141991). Stocks obtained from the Bloomington Drosophila Stock Center (NIH P40OD018537) were used in this study.

## Author contributions

This study was conceived by T.T.P, and C.C.

T.T.P performed all of the experiments with help from A.M., E.L., N.L., and J.H.

A. M also generated the CAAX::vhhGFP4 and PhyB::mcherry::CAAX constructs.

Training and access to AFM microscopy was provided by J.H and D.J.M.

T.T.P and C.C wrote the paper.

## Materials & Correspondence

Material requests and other inquiries should be directed to.

## Methods

### Fly strains and genetics

The following mutant alleles were used: *pins^P89^* (Yu et al., 2000), *pins^P62^* (Yu et al., 2000), *rod^H4.8^* (Basto et al., 2000), *sqh^AX3^* (Jordan and Karess, 1997).

Transgenes and fluorescent markers: *worGal4, pUAST-Cherry::Jupiter* (Cabernard et al., 2009), *worGal4, pUAST-Cherry::Jupiter, Sqh::GFP* (Cabernard et al., 2010), *Sqh::GFP* (Royou et al., 2002), *pUAST-Cnn::EGFP* (Megraw et al., 2002), *pUAST-pH::EGFP* (Bloomington stock center), *pUAST-attB-Caax::VhhGFP4* (this work), *pUAST-attB-PhyB::mcherry::CAAX* (this work).

### Generation of transgenic lines

#### pUAST-attB-CAAX::VhhGFP4

The coding sequence of CAAX has been synthesized as oligonucleotides. VhhGFP4 (Saerens et al., 2005) was amplified by PCR (forward primer: agggaattgggaattccgccacc ATGGATCAAGTCCAACTGGTG; reverse primer: tcttctttttacgcgt GCTGGAGACGGTGACCTG) and cloned into the pUAST-attB vector using In-Fusion technology (Takara, Clontech). The resulting construct was injected into attP flies for targeted insertion on the third chromosome (VK00020, BestGene).

#### pUAST-attB-PhyB::mcherry::CAAX

The coding sequence of PhyB::mcherry::CAAX (Buckley et al., 2016) was amplified by PCR (Forward primer: GGGAATTGGGAATTCcgccaccatggtatcaggtg; Reverse primer: acaaagatcctctagaTTAcatgataacacacttggtttttg) and cloned into the pUAST-attB vector using In-Fusion technology (Takara, Clontech). The resulting construct was injected into attP flies for targeted insertion on the third chromosome (VK00033, BestGene).

### Colcemid and Y-27632 experiments

For Y-27632 and colcemid and experiments we used *worGal4, pUAST-Cherry::Jupiter, Sqh::GFP; pUAST-PhyB::mcherry::CAAX* and *worGal4, pUAST-Cherry::Jupiter, Sqh::GFP; rod^H4.8^,* respectively. Dissected brains were incubated with Y-27632 (LC Labs) in live imaging medium at a final concentration of 62.5 *μ*gmL^−1^, or with colcemid (Sigma) at a final concentration of 25 *μ*gmL^−1^. Live imaging was acquired ~ 30 min after drugs addition. Complete spindle depolymerization was seen at the start of imaging for colcemid addition. Significant reduction of Sqh::EGFP on the neuroblast cortex was also seen ~ 30 minutes after Y-27632 addition.

### Live Cell Imaging

For all live cell imaging experiments third instar larvae were used. The live imaging procedure was performed as described previously (Cabernard and Doe, 2013) with the following minor modifications: S2 Media (Invitrogen) was supplemented with 10% HyClone Bovine Growth Serum (BGS, Thermo Scientific SH3054102). The larval brains were dissected in the supplemented S2 media and transferred into a μ-slide angiogenesis (ibidi). Live samples were imaged with an Andor revolution spinning disc confocal system, consisting of a Yokogawa CSU-X1 spinning disk unit and two Andor iXon3 DU-897-BV EMCCD cameras. A 60X/1.4NA oil immersion objective mounted on a Nikon Eclipse Ti microscope was used for most images. Live imaging voxels sizes are 0.22 × 0.22 × 1*μ*m (60x/1.4NA spinning disc).

*pUAST-CAAX::VhhGFP4* larvae were kept at 18 °C and then incubated at 29°C up to 6h prior to imaging.

### Image Processing and Measurements

Live cell imaging data was processed using Imaris x64 7.5.4 and ImageJ. Andor IQ2 files were converted into Imaris files using a custom-made Matlab code. Average intensity projections were generated in ImageJ. All Kymographs obtained from a line drawn from the apical to the basal cortex were generated with a 5 pixel wide. All Kymographs obtained from a line drawn across the furrow position site were generated with a 9 pixel wide. The intensity values at the apical, basal and furrow site cortex, as well as in the cytoplasm were extracted using a custom-made Matlab code. For the intensity plots, the cortical intensities were normalized against the cytoplasmic values. The intensity of furrow site cortex was obtained from an average value between the left and the right furrow cortex values. The curvature analysis was performed in ImageJ, using a custom made Matlab code. Cortical intensities along the cell boundary were obtained from an average intensity of three closest pixels located perpendicular to the curve.

Figures were assembled using Adobe Illustrator CS6 and all quantifications were performed in Matlab and Microsoft Excel.

### Primary larval neuroblast culture procedure

Primary neuroblast cultures were prepared as previously reported (Berger et al., 2012). In brief, wild type brains were dissected in Chan and Gehring’s medium and incubated with Collagenase Type I (Sigma) and Papain (Sigma) at 30° C for 30 min at a final concentration of 1mg/mL The brains were gently washed twice with 400 *μ*L of supplemented Schneider’s medium. The brains were then placed in a 1.5 mL tube with 200 *μ*L of supplemented Schneider’s medium and homogenized using a 200 *μ*L pipette tip by pipetting several times until the solution looked homogenous.

### Curvature analysis

To determine the local curvature along the cell cortex, a line was manually drawn in ImageJ from the apical to the basal cortex on the mid-plane. Local cortical curvature *K* can be determined via the following formula: 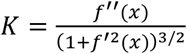, where *x* and *f(x)* are the horizontal and vertical position of the drawn cortex, respectively. The first and second derivatives (*f′(x)* and *f″(x)*) of the curve were calculated numerically using second order difference methods. Custom-written Matlab codes were used to determine curvature values for all points on the curve.

### Correlation plots

To determine the correlation between any two quantities of interest which were acquired independently, a normalized root-mean-square deviation method was used, hereby defined as the deviation coefficient. To obtain the deviation coefficient for all regions at a particular time point, the two curves were first normalized before the absolute difference was calculated. The deviation coefficients for all time point throughout mitosis was accumulated and normalized to their maximum value (NRMSD). Thus a low deviation coefficient would indicate a high correlation value due to low difference in value and vice versa.

### Kymograph quantification

Cell mid-planes were first generated using the Oblique Slicer tool in Imaris (Bitplane) and the entire image volume was then resliced along the direction of this plane for all time points. Using ImageJ, an average intensity projection was generated from three selected planes closest to the mid-plane. This procedure was done for all acquired time points. Kymographs were then generated by drawing a five pixel wide straight line from the apical to the basal cortex for all time points. To determine cortical intensity signal from a kymograph, a spline curve was drawn along the cortex on the kymograph and the XY coordinates of this curve were exported to a text file. Custom made Matlab codes were written to extract the exact XY coordinates of the drawn curve from the text file without any repetitive time points by using a standard interpolation method. Intensity signal of the drawn curve was calculated from the grayscale kymograph image using an average intensity of the three pixels, closest to the curve.

### Onset of expansion/constriction quantifications

The position of the apical, basal and furrow site cortices were traced by drawing a spline curve along each of these boundaries using the Kymographs generated earlier. The onset of expansion was chosen at the time point when the cortex expands to 3% of the cell metaphase radius (i.e. radius at 30s before anaphase onset was used in this case). The onset of constriction was set in a similar manner. This 3% change in radius value was chosen because it is equivalent to a one pixel change on the image obtained from the 60X objective, which can also be detected by the naked eye.

### AFM measurements

All stiffness measurements were performed using a Nanowizard II JPK AFM (JPK Instruments, Germany), coupled with an inverted optical microscope (Zeiss Axiovert 200, Germany). All experiments were performed in solution using a contact mode. In order to measure cortical stiffness, isolated wild type neuroblasts with fluorescent markers *Sqh::EGFP* and *Cnn::EGFP* were plated on a WPI glass bottom disc. The WPI disc was coated with 10 *μ*gmL^−1^ of Concanavalin A to anchor the cell to the glass surface to prevent slippage when pressing the cell with the AFM tip. The sample was placed on a JPK stage which was mounted on top of a Zeiss epi-fluorescent microscope. A Plan Apochromat 63X/1.4NA oil immersion objective was used together with the Zeiss AxioCam MRm monochrome digital camera to sequentially acquire fluorescent images with cortical stiffness measurements. Live imaging voxels sizes are 0. 0993 × 0. 0993 × 1 *μ*m (63x/1.4NA). High aspect ratio bead tip with a nominal spring constant 0.2 N/m and a tip height in the range 0f 10-15 *μ*m with a spherical end of radius 300 nm (B300_CONTR, Nanoandmore, Germany) was employed. A force-volume line scan consists of 30 equally spaced measurement points along 20 *μ*m distance was used to probe local cortical stiffness with a maximal applied force of 0.6 nN. The ramp velocity and the extension length were set at 20 *μ*m/s and 8 *μ*m, respectively. Each image stack and the subsequent line scan took approximately 30s to complete.

### Cortical stiffness quantifications

Cortical stiffness is obtained from the slope of the linear part of the force-depth curve. Data with a force range from 60% to 100% of the maximal force was used to fit a linear regression curve (see SF. 2a). To determine the reliability of the cortical stiffness extracted from the fitted line, the mean square error of the fit was used. The extracted cortical stiffness is classified as a reliable measurement when the mean square error of the fit is lower than the mean peak-peak fluctuation of the system, which is ~0.015 nN. Only reliable measurements were used to determine statistical average for each local region along the cell.

In order to determine the location of the AFM measurement points relative to the cell spindle-axis, precise location of the cell centrosomes and the contact point where the AFM tip touches the cell’s surface is required. The contact height for any measurement point was determined from the point along the force vs indentation height curve that fit the curve best with two linear lines (see SF. 2b). The position of each centrosome was obtained from the xyz location of a spot placing at the highest intensity signal within the centrosomal marker, Cnn::EGFP, using Imaris software. The xy position of all AFM measurement points were determined by recording the first and the last position of the AFM tip. The centrosome height and the AFM contact height were synchronized using the cell midplane at metaphase as the reference height. Location of the cortical stiffness relative to the spindle axis between the two centrosomes was determined using vector projection (see SF. 2c). At any cell cycle stage, the distance between the two centrosomes was binned into 20 equally distributed regions and reliable cortical stiffness measurements were accumulated for statistical average.

### Pressure quantifications

Isolated wild type neuroblasts with fluorescent markers *pUAST-Cherry::Jupiter* and *pUAST-PH::EGFP* were plated on a WPI glass bottom disc without any coating. The sample was placed on a JPK Cellhesion 200 stage which was mounted on top of a Zeiss confocal microscope (LSM 700). A custom-made flat PDMS wedge tip were held constantly at 5 *μ*m above the glass bottom surface where an interphase neuroblast sit. As the neuroblast enters mitosis, it rounds up and exerts a force on the wedge AFM tip, which can be detected by the photodetector (see Sup Fig. 2d). A Plan Apochromat 63X/1.4NA oil immersion objective was used to simultaneously image the neuroblast at every 1 min while recording the exerted force. Hydrostatic pressure can be determined by taking the ratio of the detected forces and the area that was in contacted with the wedge (Stewart et al., 2013). Contact area calculated from uniform tension model was used to correct the overestimated contact area detected from the fluorescent signal at the contact surface due to large point spread function in the z-axis.

---

**Supplemental Figure 1: Cultured neuroblasts can be used to measure biophysical parameters such as cortical tension and pressure.**

(**A**) Representative image sequence of a cultured wild type neuroblast expressing the apical polarity marker Baz::GFP (green; top row), the basal cell fate determinant Miranda (red; middle row) and the cell cortex marker Sqh::GFP (green; bottom row). All shown cells co-express the mitotic spindle marker pUAST-cherry::Jupiter (white). Scatter plots showing (**B**) mitosis time and (**C**) the cell cycle time for cultured larval neuroblasts (green dots; n = 14 and 49) compared to larval neuroblasts in intact brains (blue dots; n = 26 and 17). (**D**) Graph showing the rate of change in cell length for several wild type neuroblasts in an intact brain (n=13). (**E**) Scatter plot showing the time when the peak rate of change in cell length occurs relative to anaphase onset. (**F**) Representative graph showing the raw AFM data for force versus indented height. Reliable cortical stiffness was extracted from the force vs indented height curve that fit well to a linear line for a portion of the raw data. (**G**) Representative graph showing how contact height was determined. (**H**) Schematic diagram showing how an AFM contact point can be projected onto the spindle axis to be sorted into regions for statistical averaging. Scale bar: 5 *μ*m. Asterisk denote statistical significance, derived from unpaired t-tests: *; p ≤ 0.05, **; p ≤ 0.01, ***; p ≤ 0.001, ****; P ≤ 0.0001, n.s.; not significant.

**Supplemental Figure 2: Myosin’s relocalization is essential for cortical expansion**

Representative image sequences and kymographs obtained along the apical-basal axis and the furrow region of third instar neuroblasts expressing Sqh::GFP for (**A, B**) wild type, (**C, D**) *pins* mutant, Y-27632 treated wild type neuroblasts resulting in complete (**E, F**) or partial (**G, H**) Rok inhibition, (**I, J**) pUAST-CAAX::vhhGFP expressing wild type neuroblasts and (**K, L**) colcemid treated, *rod* mutant neuroblasts. Time: seconds (s). Scale bar: 5 *μ*m.

